# Bottom-up inputs are required for the establishment of top-down connectivity onto cortical layer 1 neurogliaform cells

**DOI:** 10.1101/2021.01.08.425944

**Authors:** Leena A Ibrahim, Shuhan Huang, Marian Fernandez-Otero, Mia Sherer, Spurti Vemuri, Qing Xu, Robert Machold, Bernardo Rudy, Gord Fishell

## Abstract

Higher order feedback projections to sensory cortical areas converge on layer 1 (L1), the primary site for integration of top-down information via the apical dendrites of pyramidal neurons and L1 GABAergic interneurons. Here, we investigated the contribution of early thalamic inputs onto L1 interneurons for the establishment of top-down inputs in the primary visual cortex. We find that bottom-up thalamic inputs predominate during early L1 development and preferentially target neurogliaform cells. We find that these projections are critical for the subsequent strengthening of feedback inputs from the anterior cingulate cortex. Enucleation or selective removal of thalamic afferents blocked this phenomenon. Notably, while early activation of anterior cingulate afferents resulted in a premature strengthening of these top-down inputs to neurogliaform cells, this was also dependent on thalamic inputs. Our results demonstrate that the proper establishment of top-down feedback inputs critically depends on bottom-up inputs from the thalamus during early postnatal development.

## Introduction

Our capacity to perceive and react to a rapidly changing world relies upon the ability of the neocortex to respond accurately and precisely to sensory stimuli in a dynamic environment. Processing sensory information such as vision critically depends on our ability to attend to stimuli, gate distractors, and accurately make predictions about our surroundings (Desimone and Duncan 1995, Rao and Ballard 1999, Kastner and Ungerleider 2000, Ardid, Wang et al. 2007, Squire, Noudoost et al. 2013, Zhang, Xu et al. 2014, Huda, Sipe et al. 2020). The integration of top-down feedback from higher order brain areas with incoming sensory signals allows for such functions. Increasing evidence has implicated inhibitory cortical interneurons (cINs) in this process. While parvalbumin (PV), somatostatin (SST), and vasoactive intestinal peptide (VIP) cINs have been posited to contribute to such computations (Lee, Kruglikov et al. 2013, Fu, Tucciarone et al. 2014, Zhang, Xu et al. 2014, Leinweber, Ward et al. 2017, Schneider, Sundararajan et al. 2018), the involvement of L1 cINs has been less explored. L1 contains dense axonal projections from various brain regions and is in fact the main target if cortico-cortical projections (Oh, Harris et al. 2014, Zingg, Hintiryan et al. 2014) from higher and lower cortical areas, as well as thalamocortical projections. These projections target not only the distal apical dendrites of pyramidal neurons, but also the cINs located within L1 (Ibrahim, Schuman et al. 2020). Importantly, the fact that L1 cINs have been shown to receive both thalamic (Ji, Zingg et al. 2015) and inter-cortical connections (Leinweber, Ward et al. 2017) ideally positions them to modulate incoming sensory inputs and affect excitatory responses (Zhu and Zhu 2004, Jiang, Wang et al. 2013, Lee, Wang et al. 2015, Anastasiades, Collins et al. 2020). L1 cINs have been shown to play a role in, cross-modal integration (Ibrahim, Mesik et al. 2016), interhemispheric integration (Palmer, Schulz et al. 2012), associative fear learning and plasticity (Abs, Poorthuis et al. 2018, Pardi, Vogenstahl et al. 2020) as well as sensory motor integration (Mesik, Huang et al. 2019). This makes L1 cINs a likely important target for the integration of bottom-up and top-down signaling.

A considerable breadth of evidence shows that activity in the cortex is initially dominated by bottom-up signals. The cortex only later develops recurrent activity, as a result of the establishment of functional connectivity between primary and associative cortical areas (Colonnese, Kaminska et al. 2010, Luhmann and Khazipov 2018). These findings suggest that the emergence of cortical function is a sequential process that relies on distinct epochs of activity. While it is well established that sensory experience plays a crucial role in excitatory and inhibitory neuron development (Chou, Babot et al. 2013, Pouchelon, Gambino et al. 2014, De Marco García, Priya et al. 2015), whether it is required for the establishment of feedback circuits is not known.

The two largest populations of L1 cINs are the neuron-derived neurotrophic factor (NDNF)-expressing neurogliaform (NGF) and “canopy cells”. In this paper, we explore the development of the afferent connectivity to these cell types in the primary visual cortex (V1) and reveal that they exhibit substantial differences in when they receive afferent inputs. Using monosynaptic rabies tracing we find that bottom-up thalamic inputs dominate during development, whereas top-down inputs are only progressively strengthened later. Our results show that early in development, L1 NGF cells are the main recipients of thalamic inputs. By contrast, the canopy cells in L1 only received thalamic afferents at later ages. We find that bottom-up inputs from the dorsal lateral geniculate nucleus (dLGN) onto L1 NGF cells are required for the later receipt of cortico-cortical top-down afferents from the anterior cingulate cortex (ACC), which has been implicated in visual discrimination (Zhang, Xu et al. 2014, Huda, Sipe et al. 2020) and visuo-motor mismatch responses (Attinger, Wang et al. 2017, Leinweber, Ward et al. 2017) through their feedback projections to V1. Specifically, we observed that enucleation or ablation of thalamic inputs to L1 cINs prevents the strengthening of ACC afferents onto them. Conversely, premature activation of the ACC inputs resulted in precocious development of ACC input strength onto NGF cells, a phenomenon that was critically dependent on the thalamus. Altogether these findings demonstrate the importance of bottom-up thalamic inputs on the development of top-down feedback connectivity from the ACC onto L1 NGF cINs in V1.

## Results

### Developing L1 cINs switch from receiving predominantly bottom-up thalamic to top-down cortico-cortical afferents

To map L1 cINs’ monosynaptic afferent connectivity, we used the *NDNF-dgCre* driver mouse line (Tasic, Yao et al. 2018), which labels both canopy and NGF cells, which constitute ∼70% of L1 cINs (Tasic, Yao et al. 2018, Schuman, Machold et al. 2019, Ibrahim, Schuman et al. 2020). In combination with a *Cre*-dependent AAV-helper virus and a genetically modified CVS N2c strain of the G-deleted rabies (N2c-RV) (Fig.1A), we surveyed the monosynaptic connectivity of these cells. This rabies strain has been found to be both less toxic and more efficient than the originally used B19 version (Wickersham, Lyon et al. 2007, Reardon, Murray et al. 2016). AAV helper provided the necessary components for the infection of EnvA-pseudotyped rabies virus (TVA receptor), as well as for its replication and transport (G protein) (Fig. 1A). All the required helper components (i.e., N2C-G protein, TVA receptor and a eGFP reporter) were contained within a single AAV virus (Pouchelon 2020). AAV-helper and N2C-RV-mCherry (RV) viruses were co-injected in V1. As expected, the starter cells (eGFP and mCherry co-expressing cells) were primarily localized within L1(Fig.1B). We used this approach at both developmental and adult timepoints to compare the monosynaptic afferents onto L1 cINs in V1.

**Fig 1:**
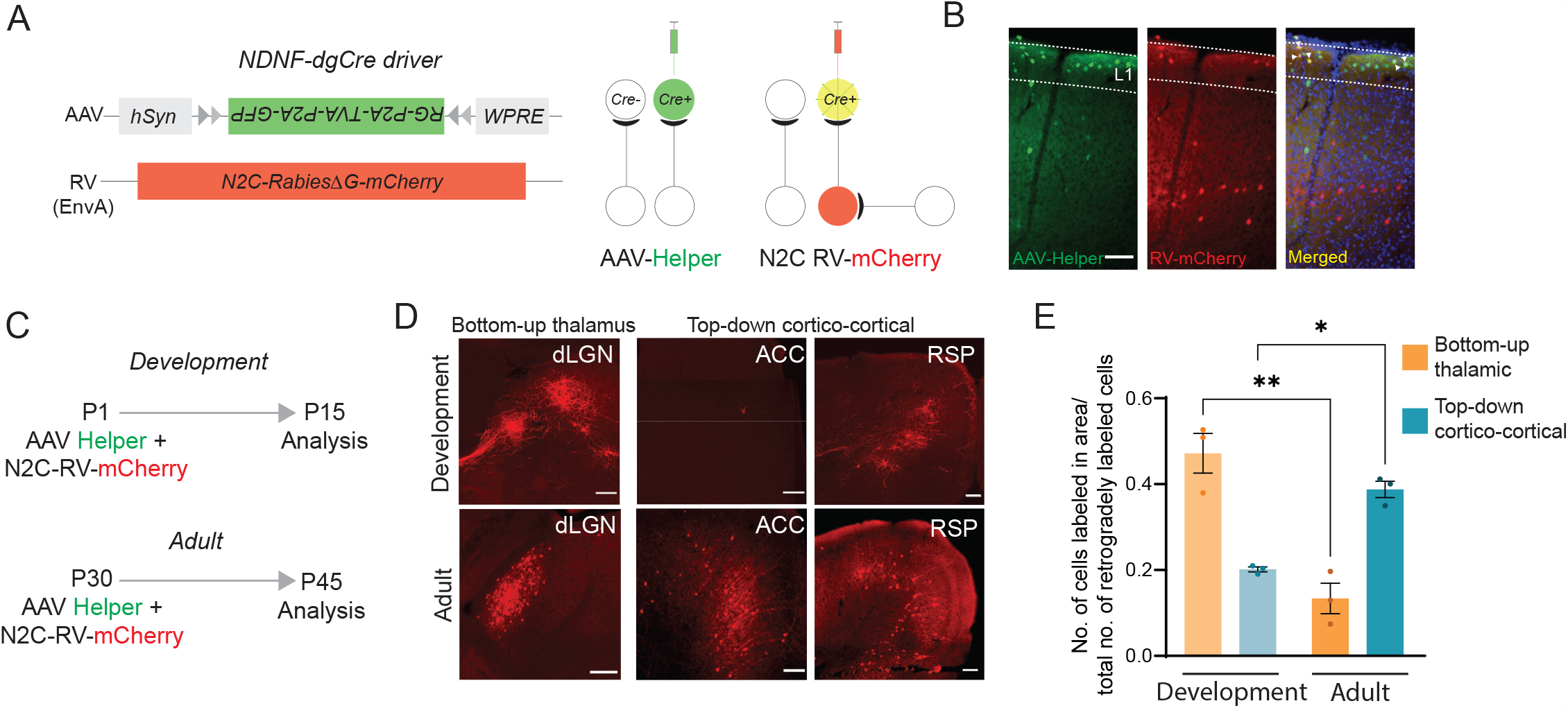
Developing L1 cINs switch from receiving predominantly bottom-up thalamic afferents to top-down cortico-cortical afferents. **(A)** Schematic of the monosynaptic rabies tracing strategy. Cre-dependent AAV-helper virus containing TVA, G protein and GFP, was injected into the *NDNF-dgCre* driver line, followed by infection and retrograde labeling of an EnvA-pseudotyped CVS-N2c(ΔG) rabies virus (N2c-RV-mCherry, in red). **(B)** Example images showing AAV-helper-infected cells (EGFP, left panel), mono-synaptically traced RV-mCherry (middle panel), and starter cells (arrowheads indicate GFP^+^ and mCherry^+^ labeled neurons in L1, right panel). Scale bar = 100µm **(C)** Developmental timeline for early AAV-helper and N2c-RV-mCherry injections at two timepoints (top panel, developmental, P1 to P15; bottom panel, adult, P30 to P45). **(E)** Examples of retrogradely labeled neurons from the dLGN (bottom-up thalamic inputs), ACC and RSP (Top-down cortico-cortical inputs) at the developmental analysis timepoint, P15 (top row), and adult analysis time point, P45 (bottom row) . Scale bar = 100µm **(F)** Quantification of the proportion of cells labeled normalized to the total number of labeled neurons in the brain (bottom-up thalamic vs top-down cortico-cortical) at development and adult time points (development bottom-up vs adult bottom-up, p value = 0.0014; development top-down vs adult top-down, p value = 0.0274; 1-way ANOVA with multiple comparisons)

To assess developmental connectivity, we injected the AAV helper and N2c-RV at P1 (n=3 animals) and analyzed the retrogradely labeled cells at P15. For the adult connectivity, we injected mice at 4-5 weeks of age (n=3 animals) and analyzed the brains similarly two weeks later (Fig. 1C). By quantifying the normalized number of retrogradely labeled neurons across regions (see Methods for details), we found that during development, the most numerous inputs that L1 cINs receive were from the primary sensory thalamus, (i.e., dLGN, Fig.1D-E; 0.471± 0.0217). In contrast, in adults, while L1 cINs maintained thalamic inputs, they were collectively less numerous than the long-range cortico-cortical inputs from the ACC and the retrosplenial (RSP) cortex (Fig. 1E; bottom-up proportion: 0.133±0.016 vs top-down proportion: 0.387±0.009). We focused on these projections, as they have been shown to have abundant axonal projections in the superficial layers of V1. The increase in long-range cortico-cortical connectivity in the adult indicates that as L1 cINs mature, their afferents shift from being predominantly bottom-up to top-down. Additionally, L1 cINs in V1 received both local inputs, primarily from L5 (pyramidal neurons and presumably SST cINs, Fig. S1A, (Abs, Poorthuis et al. 2018)) in the adult; L5 and L6 (predominantly L6b) during development (Fig. S1B and see (Meng 2020), as well as long-range cortical inputs from higher visual and from primary auditory cortex. Furthermore, they received a small number of afferents from both the posterior-parietal and temporal-association areas (data not shown).

NDNF+ L1 cINs also received inputs from higher-order visual thalamic nuclei, such as the lateral-posterior (LP) nucleus and other visual thalamic nuclei such as the lateral-dorsal (LD) (Nassi and Callaway 2009, Saalmann, Pinsk et al. 2012). Interestingly, while LP is thought to be the main source of thalamic inputs to L1 in V1 (Roth, Dahmen et al. 2016), we found that the dLGN inputs were more predominant in both development and in the adult (Fig. S1C-D).

### L1 NGF cells and L4 excitatory cells share common sensory thalamic inputs

To label bottom-up thalamic afferents, we utilized the *Vipr2-Cre* driver line, which has been previously shown to be specific for the dLGN, but not LP (Zhuang 2019). *Vipr2-Cre* mice crossed with the tdTomato reporter Ai14 labels the dLGN, ventro-basal nucleus (VB) (Fig.2A), and ventral medial geniculate body (MGBv); the visual, somatosensory, and auditory sensory nuclei in the thalamus with a high degree of specificity during both development and in the adult. To determine the strength of thalamic inputs to L1 cINs in V1, we injected Cre-dependent channelrhodopsin (hChR2-eYFP) in the dLGN and as expected thalamic fibers were observed within both L4 (the main recipient of dLGN inputs) and L1 of V1 (Fig. 2B). We injected neonatal P0/P1 mouse pups, or adult mice, and after a two-week survival period, we performed whole cell voltage clamp recordings from both L1 and L4 neurons in V1 coronal slices in response to optogenetic stimulation of dLGN fibers in V1 (Fig. 2C).

**Fig 2:**
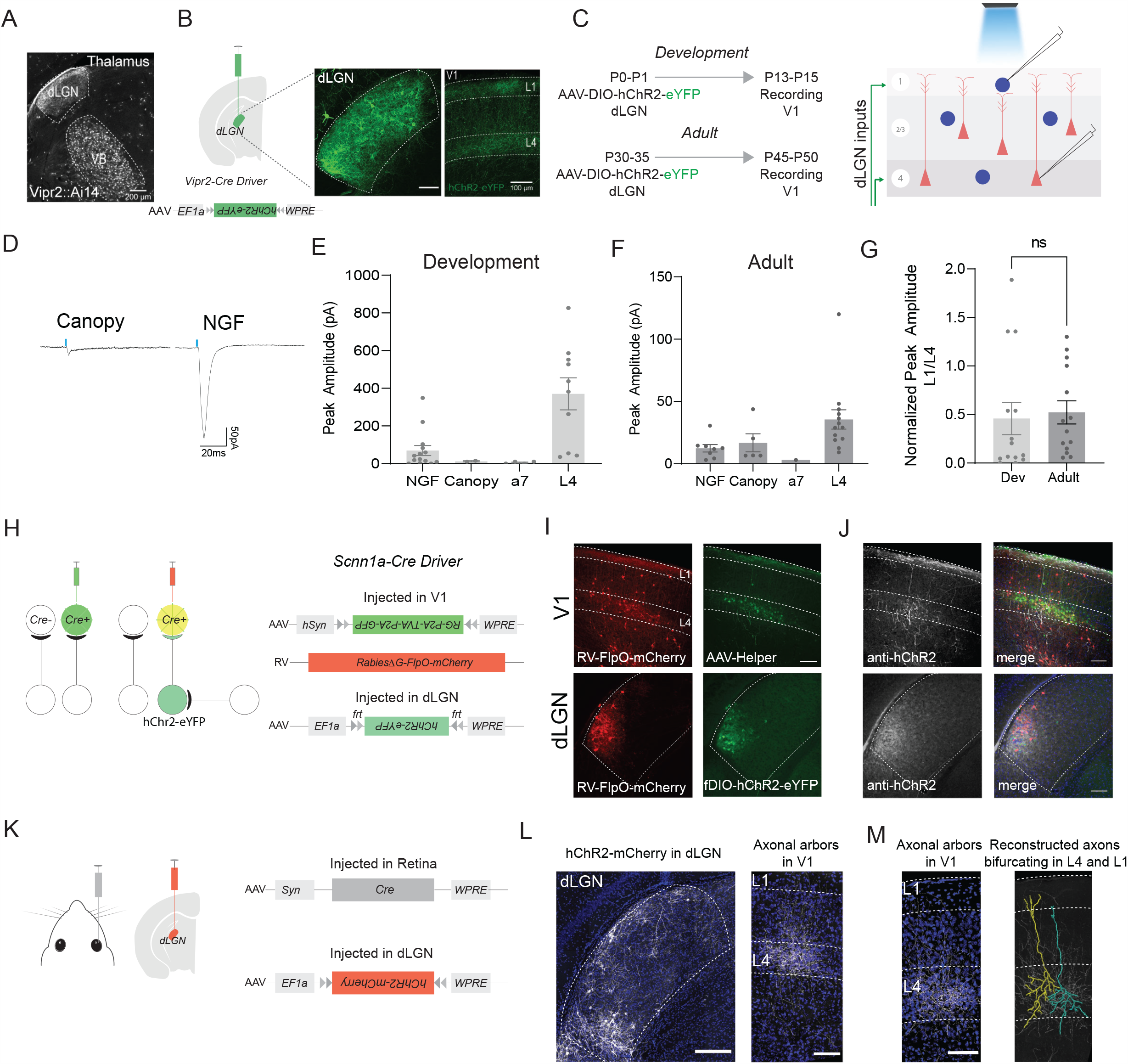
L1 NGF cells and L4 excitatory cells share common sensory thalamic inputs. **(A)** Image showing specificity for first-order thalamic nuclei dLGN and VB, using the*Vipr2-Cre* driver line crossed with Ai14 reporter line (cre-dependent tdTomato). Scale bar = 200µm. **(B)** Schematic of the AAV-DIO-hChr2-eYFP virus injection into dLGN of *Vipr2-Cre* driver line (left panel), virus expression in dLGN (middle panel) and resulting axonal projections to both L4 and L1 in V1 (right panel). Scale bar=100µm. **(C)** Left panel: Timeline for the AAV-DIO-hChr2-eYFP virus injection (P0-P1; P30-P35) and recording at development timepoint and adult timepoint (P13-P15; P45-P50). Right panel: Schematic of the slice recording to blue light stimulation from L1 cINs and L4 excitatory neurons in the same column. **(D)** Example EPSC traces to blue light stimulation of dLGN fibers in V1 L1. Canopy cell response (left panel) and NGF cell response (right panel) from a P14 *Vipr2-Cre* mouse. **(E)** Peak EPSC amplitude of all neurons recorded during development. **(F)** Peak EPSC amplitude of all neurons recorded in adult. **(G)** L1 responses normalized to L4 responses in development and adult. **(H)** Modified FlpO-expressing RV approach to test whether L4 and L1 receive common inputs. Cre-dependent AAV-helper and N2c-RV-FlpO-mCherry viruses were injected in V1 of *Scnn1a-Cre* driver line, followed by flp-dependent AAV-fDIO-hChR2-eYFP virus injection in dLGN. **(I)** Top left panel shows the N2c-RV-FlpO-mCherry expression in V1; top right panel shows the AAV-helper EGFP expression in L4 of V1. **(J)** Antibody against hChR2 labeled fibers in V1 (top left panel), dLGN (bottom left panel) and merged images from I and J in V1 (top right panel) and the dLGN (bottom right panel). Scale bar = 100µm. **(K)** Schematic to sparsely label neurons in dLGN. AAV-Syn-Cre virus was injected in the right retina results in sparsely trans-synaptic cre expression in the dLGN, followed by an AAV-DIO-hChR2-mCherry virus injection in dLGN. **(L)** Sparsely labeled neurons in dLGN (left panel), scale bar = 500µm. dLGN fibers in V1 (right panel), scale bar = 100µm. **(M)** Neurolucida tracing of dLGN axons reveals collaterals in L4 and L1.

We wanted to determine which L1 cIN subpopulation is the recipient of dLGN inputs and to what extent these inputs overlap with those onto L4. It is well known that L1 cINs are diverse. Historically they have been classified in two main groups: neurons with an axonal arbor mainly confined to L1, which include neurogliaform cells (called elongated NGFCs by (Jiang, Wang et al. 2013) because their axonal arbor spans several columns) and cells with descending collaterals to deep layers called single bouquet cells (SBCs) (Kawaguchi and Kubota 1997, Zhu and Zhu 2004, Cruikshank, Ahmed et al. 2012, Jiang, Wang et al. 2013, Muralidhar, Wang et al. 2013). Schuman et al., recently showed that the neurons that have an axonal arbor largely confined to L1 express NDNF, a marker expressed in 70% of L1 cINs. Furthermore, they showed that NDNF neurons consist of two subtypes: NGFCs and a cell type they called “canopy cells” (Schuman, Machold et al. 2019). Despite their gross morphological similarities, the two subpopulations are distinct in their expression of Neuropeptide Y (NPY), intrinsic physiological properties and output connectivity. Specifically, NGF cells are NPY^+^ (Fig S2A), late spiking and have a ramping-up membrane depolarization, with a relatively high rheobase during development (Fig S2B left panel). Canopy cells by contrast are regular spiking, display a small degree of spike adaptation and have a voltage sag in response to hyperpolarizing current injection (Fig S2B right panel).

We wished to determine whether these cell types receive thalamic inputs and if so, how the strength of these inputs compare to those targeting L4 excitatory cells (the main input layer from the thalamus). We found that NGF cells, but not canopy cells, receive considerable direct thalamic input (AMPA current recorded by clamping cells at - 70mV) during development (Fig. 2D-E, NGF cell response: 97.3±8.8pA vs Canopy cell response: 11.63±2.5pA). Interestingly, in adults the responses on canopy cells became comparable to those of NGF cells (Fig. 2F). While in both development and adults the response of L4 excitatory neurons was on average twice that of L1 cINs, in specific instances some NGF cell responses were in fact comparable (Fig. 2G, (Takesian, Bogart et al. 2018)). Notably the largest non NDNF population of L1 cINs, the alpha-7 expressing L1 cINs, were not found to receive thalamic input in either development or in the adult. Thus, across development the thalamic innervation of L1 cINs is cell type specific.

Next, we wanted to determine whether the thalamic afferents that project to L1 are shared with those that project to L4. To that end, we utilized a L4-specific *Cre* driver line (*Scnn1a-Cre*) and used a modified approach, where the RV also expresses *FlpO* recombinase together with mCherry. AAV helpers and EnvA-N2c-RV-FlpO-mCherry were co-injected in L4 of V1, followed by an injection of a flp-dependent hChr2-eYFP in the dLGN (Fig. 2H). This allowed us to examine the projection pattern of L4-projecting dLGN neurons. We used an antibody against hChR2 to visualize the fibers in V1. In addition to their projection in L4, the retrogradely labeled dLGN neurons (Fig2J lower panel) were found to have collaterals in L1 (Fig. 2J, upper panel). This demonstrates L4 and L1 share some inputs from the same thalamic axons. To independently confirm this, we sparsely labeled neurons in the dLGN, by injecting AAV virus carrying Cre recombinase in the contralateral retina (which with low probability travels anterogradely and trans-synaptically into the dLGN, (Zingg, Chou et al. 2017). This was followed by an injection of a cre-dependent hChR2-mCherry AAV in the dLGN (Fig. 2K). This allowed us to directly visualize single dLGN neurons that project to both L4 and L1 (Fig. 2L). Moreover, when the resulting axonal arborizations were traced (Fig. 2M) it revealed at the single axon level bifurcating collaterals within both L4 and L1. This further confirmed that some thalamic axons target both layers.

### ACC inputs to L1 cINs strengthen across development

To elucidate the corresponding developmental changes in the strength of top-down inputs, we focused on the feedback projections from the ACC to V1. In adults, inputs from the ACC have been shown to extensively arborize within L1, where they target various postsynaptic neurons across the different layers of V1 (Leinweber, Ward et al. 2017). However, the dynamics by which this input is established across development are unknown. To explore this question, we injected hChr2-eYFP in the ACC region (Fig. 3A) and assessed the fiber pattern (Fig. 3B) and input strength (Fig. 3C) in L1 and L2/3 of V1 both during development and in the adult. Based on our rabies tracing results, we expected to see little or no feedback projections from the ACC to V1 during development (Fig.1D-E). However, as early as P10, we observed the presence of ACC afferents distributed across the different layers of V1, which become concentrated within the superficial and the deep layers of V1 by adulthood (Fig. 3B). To assess the functional strength of these fibers during development (P13-P15) and in the adult (∼6-7 weeks), we optogenetically activated the ACC axons in V1 and recorded from the different L1 cINs subtypes, as well as L2/3 excitatory cells (Fig. 3C). To ensure that the observed differences did not result from differential levels of hChR2 expression, both ages were examined two weeks post-injection. We found that during development despite the abundance of fibers from the ACC to V1, the amplitude of responses in both L1 and L2/3 neurons were small (Fig. 3D and 3E left panel, L1 NGF response: −32.075±2,56pA; L1 Canopy response: −19.66±1.314pA; L2/3 excitatory neuron response: −7.5±0.5pA). Responses seen in L1 were still on average greater than those observed in L2/3, a trend that becomes even more pronounced in the adult (Fig.3E, right panel). Specifically, the peak amplitude of AMPA currents (recorded by clamping the cells at −70mV) increased by ∼10 folds in L1 cINs by adulthood (Fig. 3F). Interestingly, L1 NGF cells were the main recipients of these feedback inputs. By contrast, the inputs onto canopy and alpha-7 cells (data not shown) were considerably smaller and were roughly comparable to those found onto L2/3 excitatory cells (Fig. 3E, L1 NGF response: −171.6±3.03pA, L1 canopy response: −38.33±2.179pA, L2/3 excitatory neuron response: −21.75±1.03pA). These results suggest that the developmental refinement and strengthening of feedback from ACC to V1 is developmentally regulated and occurs in a progressive and cell-type specific manner.

**Fig 3.**
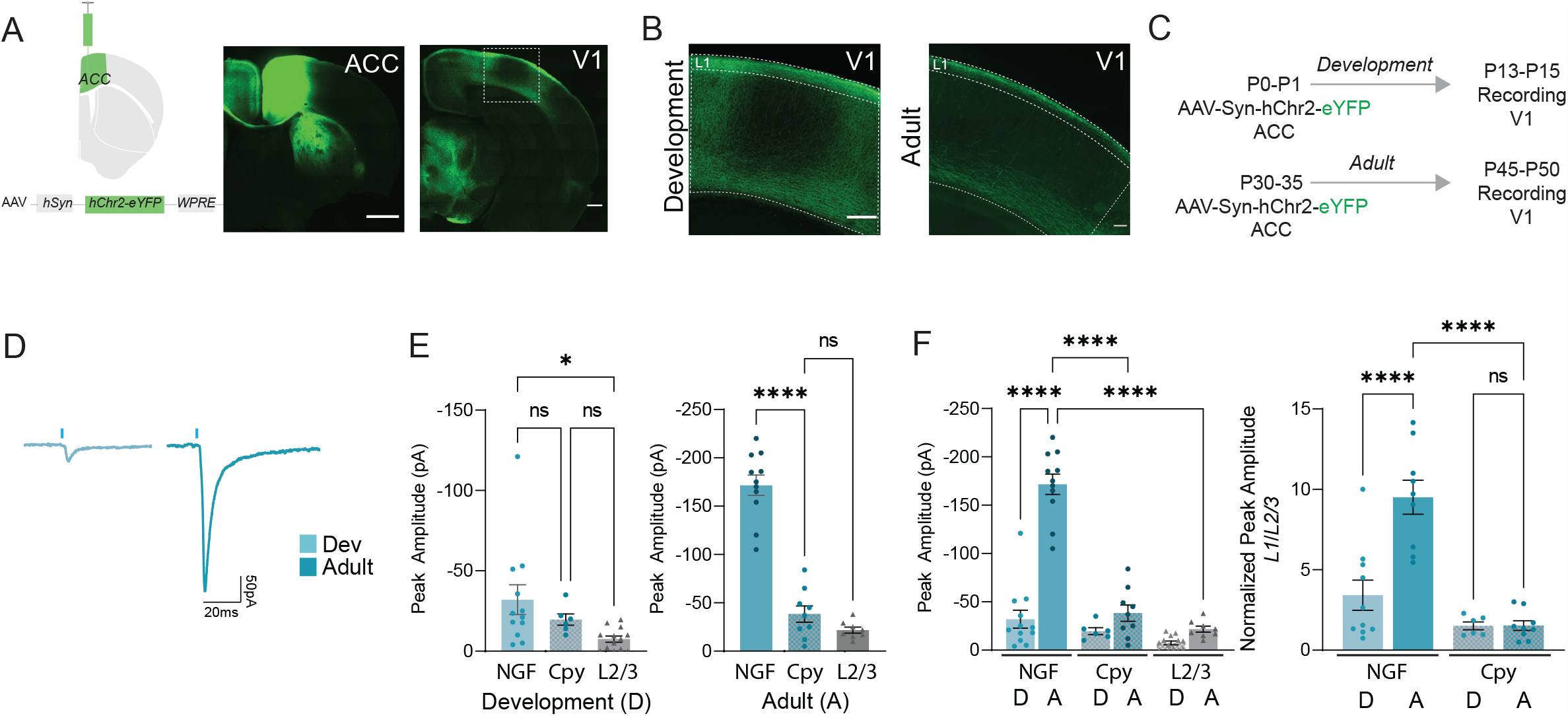
ACC inputs to L1 cINs strengthen across development. **(A)** Schematic of the hChR2-eYFP virus injection and eYFP expression in the ACC during development (left panel) and its axonal pattern in V1 (right panel). Scale bar = 500µm. **(B)** Magnified view of the boxed region in A (left panel, development), and image of the axonal pattern in the adult V1 (right panel, adult). Scale Bar=100µm. **(C)** Timeline of virus injection in ACC and recording in V1 for both development and adult timepoints. **(D)** Example traces of a NGF cell in V1 response to optogenetic stimulation of ACC fibers in development (left) and adult (right). **(E)** Peak EPSC amplitude of L1 NGF and canopy (Cpy) cells and L2/3 pyramidal neurons during development (left) and in the adult (right). One-way ANOVA with multiple comparisons. *=<0.05; ***=<0.0001. **(F)** Pair-wise Comparison of the same data in E in development (D) and adult (A) (left panel) and same comparison after normalizing to L2/3 pyramidal neurons in the same slice (right) ***=<0.0005 ****=<0.0001 unpaired t-test

### L1 NGF cells require bottom-up sensory inputs for the strengthening of top-down connections

The observation that bottom-up sensory afferents onto L1 cINs are established prior to their receipt of ACC inputs prompted us to ask whether the developmental refinement of feedback inputs is dependent on sensory activity. To do so, we examined the impact of sensory deprivation on ACC afferent development by early postnatal enucleation of the eyes(Fig. 4A). We found that in V1, this resulted in a significant decrease in the strength of ACC inputs onto L1 NGF cells (Fig. 4B). Interestingly, this did not result in any drastic morphological changes in L1 cINs (data not shown). This indicates that thalamic activity is required for the strengthening of feedback inputs to the visual cortex. In addition, this also results in an increase in the frequency of spontaneous synaptic events onto L1 cINs, perhaps as a consequence of a compensatory increase in total excitatory inputs stemming from the enucleation (Fig.S3A-C) as seen in other cell types in the cortex (De Marco García, Priya et al. 2015, Frangeul, Kehayas et al. 2017).

**Fig 4.**
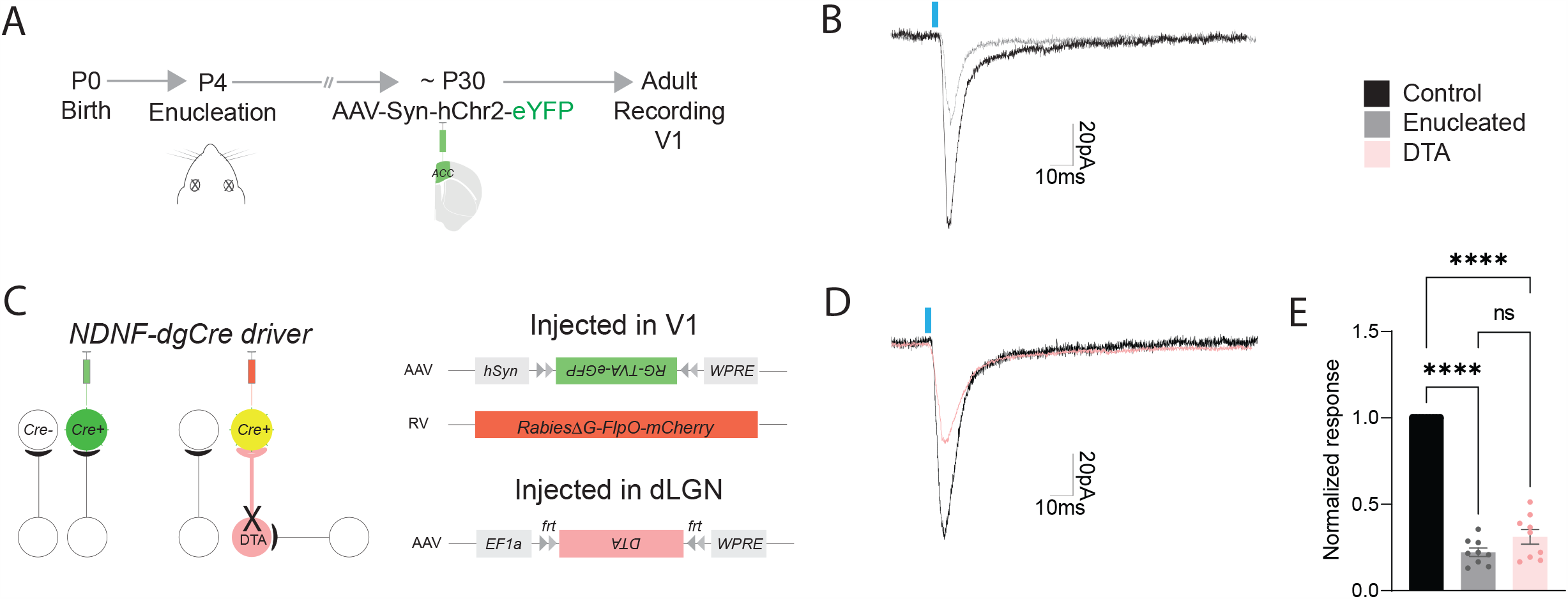
L1 NGF cells require bottom-up sensory inputs for the strengthening of top-down connections. **(A)** Timeline of enucleation (P4), Injection of AAV-hChr2 (∼P30) into ACC and recording in L1 cINs in V1 in adults (∼P45). **(B)** Example traces of a NGF cell in V1 response to optogenetic stimulation of ACC fibers in control (black) and enucleated (grey) animals. **(C)** Schematic of the FlpO-dependent DTA ablation of L1 projecting dLGN axons. Cre-dependent AAV-helper and N2C-RV-FlpO-mCherry viruses injected in V1 of Ndnf-dgCre mouse; *Flp*-dependent AAV-fDIO-DTA injected in the dLGN. The timeline for DTA ablations was similar to that in A. **(D)** Example traces of a NGF cell in V1 response to optogenetic stimulation of ACC fibers in in control (black) and DTA-dLGN ablated afferents (orange) animals. **(E)** The normalized responses of enucleated and DTA experimental groups compared to wild type controls. ****=<0.0001 unpaired t-test.

As enucleation must result in considerable indirect effects through its impact on other areas of the visual system (Williams, Reese et al. 2002, Nahmani and Turrigiano 2014, Rose, Jaepel et al. 2016, Jaepel, Hübener et al. 2017, Bhandari 2020, Hooks and Chen 2020), we wished to see if selective manipulation of the thalamic inputs that specifically project to L1 result in a similar phenotype. To do so, we injected AAV helpers together with RV-FlpO-mCherry virus in *NDNF-dgCre* pups and followed that with an injection of a *Flp*-dependent diphtheria toxin subunit A (DTA) in the dLGN (Fig. 4C). This allowed us to selectively ablate only those dLGN neurons that innervate L1 cINs (Fig. S3D), with the caveat that the common inputs between L4 and L1 will also be disrupted. Importantly the RV-infected L1 cINs remain physiologically healthy up to 5 weeks post rabies infection (data not shown and see (Reardon, Murray et al. 2016). Upon reaching adulthood, hChR2-eYFP virus was injected into the ACC and following a two-week survival period the RV-injected starter cells were recorded in V1. Similar to what was observed in the enucleation experiments, we found a significant decrease in the strength of the ACC inputs onto L1 cINs (Fig. 4D-E). Notably, in enucleated animals, a similar reduction was not observed within L1 cINs in A1 cortex (data not shown). These results suggest that bottom-up sensory inputs are required for the strengthening of feedback projections from the ACC onto L1 cINs in V1.

### Top-down connectivity onto L1 NGF cells depends upon coordination with bottom-up inputs

Given that compromising bottom-up sensory signaling prevents the strengthening of ACC inputs onto L1 NGF cells in V1, we wondered if early activation of either dLGN or ACC inputs would result in the premature strengthening of the latter. To explore this question, in separate cohorts of pups, we either injected the dLGN of *Vipr2-Cre* mice (∼P1) with a DIO-hM3D(Gq)-DREADD-mCherry AAV or the ACC with a hM3D(Gq)--mCherry-expressing AAV (Fig. 5A-B). In both circumstances, to measure the strength of top-down inputs, we also injected hChR2-eYFP in the ACC. In the case of ACC-Gq-DREADD activation, we found that in the ACC, at least 70% of the neurons expressed both mCherry and eYFP, as well as responded to both optogenetic and chemogenetic activation (Fig S4A-C).

**Fig 5.**
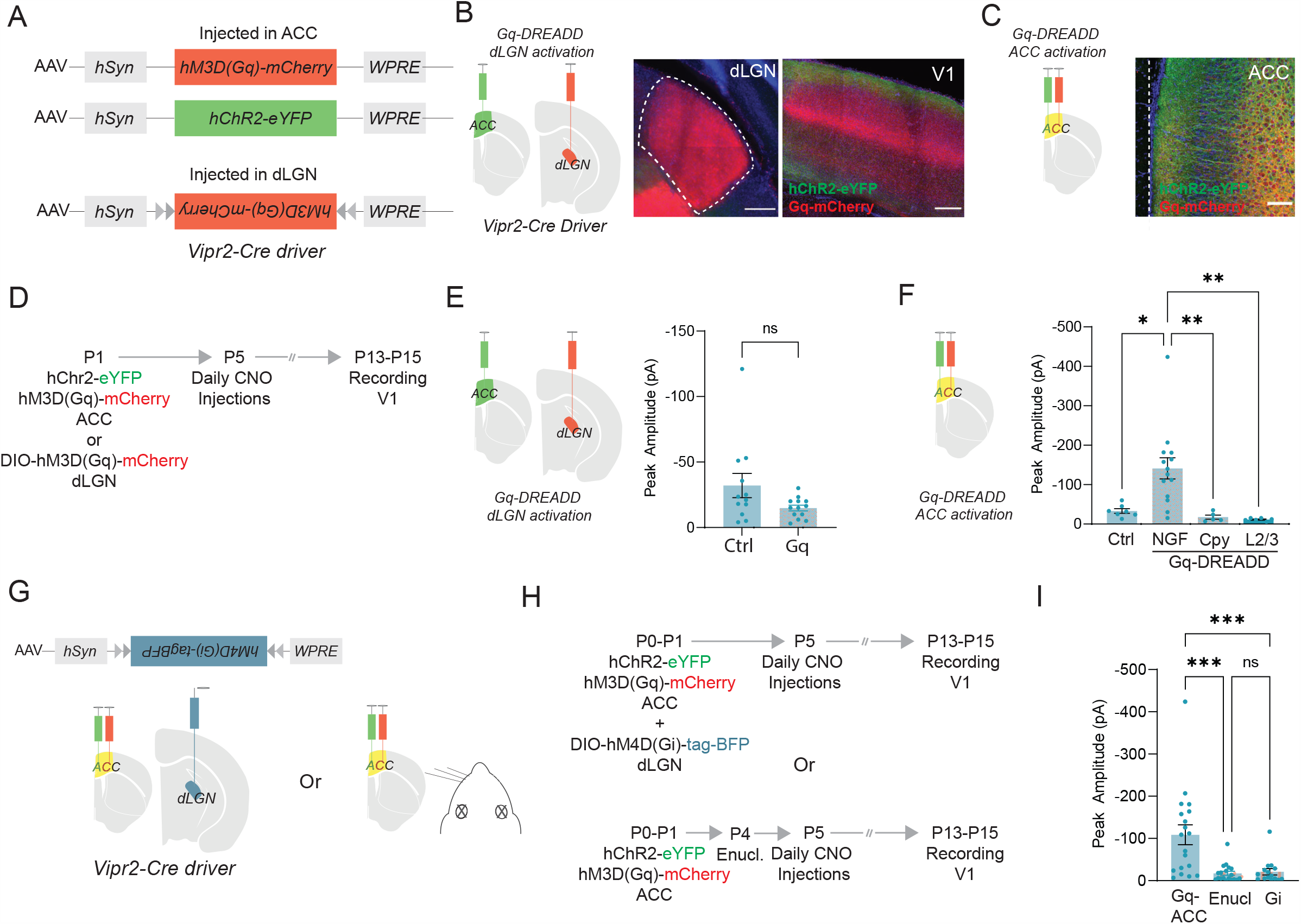
Top-down connectivity onto L1 NGF cells depends upon coordination with bottom-up inputs. **(A)** AAVs expressing either hM3D(Gq)-DREADD-mCherry and hChR2-eYFP were co-injected in the ACC Or AAV-DIO-hM3D(Gq)-DREADD and hChR2-eYFP in the ACC were co-injected in the dLGN of Vipr2-cre mice (as appropriate). **(B)** Schematic of the Gq-DREADD dLGN-activation experiment (left panel), the expression of Gq-DREADD-mCherry in the thalamus (middle panel) and co-expression of Gq-DREADD-mCherry in the dLGN and hChR2-eYFP axons in V1 (right panel). Scale bar = 200µm. **(C)** Schematic of the Gq-DREADD ACC-activation experiment (left panel) and co-expression of Gq-DREADD-mCherry and hChr2-eYFP in ACC (right panel). Scale bar = 100µm. **(D)** Timeline of AAV virus injection (P1), CNO administration (daily starting at P5) and recording of L1 cINs in V1 (∼P13-P15). **(E)** Comparison of peak EPSC amplitude response of L1 cINs (both NGF and canopy cells) to ACC hChR2 stimulation: control vs Gq-DREADD dLGN activation. ns=>0.05 **(F)** Comparison of peak EPSC amplitude responses of the different cell types in V1 to ACC hChR2 stimulation in control vs Gq-DREADD ACC activation. Cpy=Canopy Cells. One-way ANOVA with multiple comparisons *=<0.05; **=<0.01. ns comparisons not shown **(G)** Schematic of the hM4D(Gi)-DREADD dLGN-inhibition experiment, while simultaneously activating ACC with Gq-DREADD (left panel) and schematic of the enucleation combined with activation of ACC with Gq DREADD (right panels). In both experiments, ACC was injected with AAV-hChR2 as in E-F. **(H)** Timeline of AAV virus injections (P1), CNO administration (daily starting at P5) recording in L1 cINs in V1 (∼P13-P15, top panel) or enucleation timeline (bottom panel). **(I)** Comparison of peak amplitude response of L1 NGF cells to hChR2 stimulation: Gq-DREADD ACC Activation condition (labeled as Gq-ACC, left bar) vs Gq-DREADD ACC activation + enucleation condition (labeled as Enucl, middle bar) and Gq-DREADD ACC activation together with Gi-DREADD dLGN inhibition condition (labeled as Gi, right bar). One-Way ANOVA with multiple comparisons ***=<0.005

In these two experimental groups, we then compared the strength of ACC projections onto L1 cINs at ∼P14 following daily injections of Clozapine-N-oxide (CNO) or saline (starting at P5, Fig. 5C). We did not observe strengthening of ACC projections when the dLGN pathway was over-activated (Fig. 5E). In contrast, selective activation of the ACC pathway resulted in a significant increase in the ACC input strength onto L1 cINs (Fig. 5F). Moreover, this increase was specific to the NGF cell population, and was not observed in canopy, a7 or L2/3 excitatory neurons (Control responses to ACC activation: −35.5±2.03pA vs Gq-DREADD-ACC NGF cell response: −163.529±6.84pA, Gq-DREADD-ACC Canopy cell response: −17.4±2.08pA, Gq-DREADD-ACC L2/3 response: −11.4±0.55pA). While the premature increase in strength could be attributed to a general plasticity mechanism, the specificity by which this occurred was very striking.

We wondered whether the augmentation in the strength of ACC afferents to L1 NGF cells upon increased activation of ACC projections was dependent upon bottom-up sensory thalamic signals. We reasoned that this may occur due to the coordinated engagement of bottom-up and ACC afferents impinging onto L1 NGF cells (Fig. 4). We therefore hypothesized that this augmentation should be dependent upon bottom-up sensory activity. We therefore tested whether dampening of sensory signaling could impair this phenomenon. To achieve this, we either injected a AAV-DIO-hM4D(Gi)-DREADD into the dLGN (in *Vipr2-Cre* pups) or we enucleated the mice at P4, while expressing Gq-DREADD and hChR2 in ACC (Fig. 5G-H). Consistent with our hypothesis, both manipulations reversed the ability of Gq-DREADD activation of ACC from increasing the synaptic strength of ACC afferents onto L1 NGF cells (Fig. 5I).

Next, we queried whether there exists a threshold requirement for bottom-up activity or if enhanced co-activation of ACC and dLGN afferents would further increase the strength of the ACC inputs. To explore this question, we injected the dLGN of *Vipr2-Cre* pups (∼P1) with a DIO-HM4D(Gq)-DREADD-mCherry AAV and the ACC with a HM4D(Gq)-DREADD-mCherry-expressing AAV and as in earlier experiments co-injected hChR2-eYFP in the ACC (Fig. S4D, left panel). As expected, upon CNO administration the strength of ACC afferents was increased (Fig. S4D, right panel). However, this did not increase the strength of the ACC afferents beyond what we observed when we activated them in the absence of enhanced dLGN activation (Fig. S4E). We conclude that while activity of dLGN is required for the enhancement of ACC connectivity, additional activation of dLGN does not result in further strengthening of ACC afferents.

## Discussion

Interneurons in superficial visual cortex have been shown to receive both bottom-up sensory inputs and top-down cortico-cortical afferents from ACC. Despite the prevalence of ACC fibers in L1 of V1, previous studies have not explored the extent to which cINs in this layer receive these inputs. Here, we observe that among L1 cINs, NGF cells are notable in that they receive these inputs, as well as bottom-up sensory afferents. In addition, we demonstrate that a portion of the dLGN afferents to L1 are shared with those in L4. Furthermore, we find that during development, the strengthening of ACC afferents onto L1 NGF cells requires earlier receipt of thalamic inputs from the dLGN. In exploring the role of coordinated activity between ACC and thalamic fibers, we find that ACC afferent activity can modulate the strength of these top-down inputs. By contrast, while thalamic afferents are necessary for acquisition of ACC inputs, a threshold level of activity suffices.

### L1 NGF cells receive early thalamic inputs

In our rabies tracing of L1 cINs in V1, we observed that thalamic afferents originate from both dLGN, as well as higher-order LP thalamus. While dLGN is thought to convey primarily sensory signals, the LP receives input from a variety of sources, including the superior colliculus (Gale and Murphy 2014, Fang, Chou et al. 2020) as well as cortical feedback (Bennett, Gale et al. 2019). This suggests that the activity in LP reflects higher order signaling from a variety of modalities in addition to visual inputs. How these two sources of thalamic input are coordinated within L1 cINs is poorly understood. Recent work has implicated the LP in functioning to suppress noise in the visual cortex through a L1 mechanism (Fang, Chou et al. 2020), however the specific cINs within L1 that mediate this signaling remain unknown. Despite the presence of both first- and higher-order thalamic inputs to NDNF^+^ L1 cINs, our findings indicate that in this layer, these INs are more abundantly innervated by the dLGN (Fig. S1C-D).

Previous work from our laboratory and others have demonstrated that superficial NGF cells receive thalamic inputs early (De Marco García, Priya et al. 2015, Che, Babij et al. 2018). Despite their positioning in superficial cortex, by P15 NGF cells are receive bottom-up activity. They perhaps receive these afferents as early as the SST cINs in deep layers, the latter of which are the first born cIN population (Marques-Smith, Lyngholm et al. 2016, Tuncdemir, Wamsley et al. 2016, Pouchelon 2020). Given the efferent copy of first-order thalamic inputs to L4 and L1 in V1, it is appealing to consider whether the coordinated induction of excitation within L4 and inhibition within L1 may have developmental functional significance. Nonetheless, despite the considerable strength of dLGN inputs onto L1 NGF cells (Fig. 2E), our preliminary examination of their output during development (results not shown) suggests that these connections do not produce strong inhibition. Instead, our results indicate that the sensory thalamic inputs to L1 NGF cells are essential for the promotion of afferent connectivity from the ACC, perhaps through a coincident depolarization and Hebbian mechanism. Interestingly, previous work from our group indicates that early thalamic inputs to S1 function competitively with local excitatory inputs to NGF cells (De Marco García, Priya et al. 2015). Specifically, when NMDA signaling, which we previously posited disproportionately drives thalamic versus cortical inputs onto NGF cells, is ablated, it results in a shift in the connectivity of these cells from receiving thalamic to local cortical afferents. Notably, this study focused on L2/3 NGF cells rather than those in L1. Our present work reveals that thalamic afferents onto L1 NGF cells in visual cortex function cooperatively with long-range inputs from ACC, resulting in their developmental strengthening. Taken together, these results indicate that local and long-range connectivity onto L1 NGF cells are centrally dependent on early dLGN afferents.

### L1 NGF cells function in the coordination of bottom-up and top-down signaling

While L1 cINs have been previously shown to receive bottom-up sensory inputs (Ji, Zingg et al. 2015), we demonstrate that this is specific to the NGF cell population during development. In addition, the dLGN afferents to L1 NGF cells at least in part represent efferent copy of the sensory afferents to the L4 cells. This suggests that NGF cells in L1 are uniquely positioned to coordinate the activity of bottom-up and top-down signaling in the superficial layers of the visual cortex. It is likely that thalamic first-order afferents to the NGF cells synergize with projections from other sources. While previous studies implicate that cholinergic afferents may subserve this role (Letzkus, Wolff et al. 2011, Alitto and Dan 2012, Fan, Kheifets et al. 2020), our work implicates the co-recruitment of ACC afferents as a potential source for their engagement.

L1 NGF cells have late-spiking properties and are thought to signal through volume transmission, which provides strong and sustained inhibition in superficial visual cortex. This positions them to function in suppressing activity when bottom-up and top-down signals are coordinated. This could be potentially useful in contexts where the expected internal representation is in concordance with bottom-up sensory signals, indicative that sensory expectations are met. They therefore could be functioning in a manner akin to what has been proposed for PV interneurons, which have been implicated in the suppression of self-generated movements in the auditory cortex (Schneider, Sundararajan et al. 2018). In complementary fashion to the L1 NGF cells, VIP cINs in L2/3 also receive top-down signals including from ACC (Zhang, Xu et al. 2014), but notably not direct bottom-up inputs (Ji, Zingg et al. 2015). VIP cINs disinhibit SST cINs (Lee, Kruglikov et al. 2013, Fu, Tucciarone et al. 2014), the latter of which has been shown to be activated during visual-learning (Attinger, Wang et al. 2017). Thus, emerging evidence has begun the reveal how different cIN populations are coordinated to support cortical computations. Of the major cIN classes, the L1 NGF cell has been the least studied. Our findings provide insight as to the contexts in which they are likely recruited. Interestingly, we have observed that the NGF cells in the auditory cortex are organized similarly to those we report here (data not shown). We are currently exploring how these populations are engaged *in vivo* in both auditory and visual tasks.

## Methods

### Mice

All experiments were approved by and in accordance with Harvard Medical School IACUC protocol number IS00001269. C57Bl/6 mice were used for breeding with transgenic mice. Transgenic mice, NDNF-dgCre (stock number: 028536), Vipr2-IRES-Cre-D (stock number: 031332), Scnn1a-Tg3-Cre (stock number: 009613), Ai14 (expressing tdTomato, stock number: 007909) are available at Jackson Laboratories. Both female and male were used in the entire study.

### Sensory deprivations

To deprive mice from visual sensory input enucleation was performed. P4 mouse pups were anesthetized by hypothermia. A small incision was made between the eyelids with a scalpel and the eye was separated from the optic nerve with micro-scissors to be removed from the orbit. The incision was secured using biocompatible Vetbond glue. The pups were then allowed to recover on a heating pad before being returned to their mother.

### Histology

Mice at between P42-P46 for the adult time point or P15 for the developmental time point were transcardially perfused with 4% paraformaldehyde (PFA) and brains were fixed overnight in 4% PFA at 4 °C. 50 μm vibratome sections were used for all histological experiments. Every 3rd representative section was collected, and the sections were processed for immunohistochemistry. For immunofluorescence, brain sections were incubated for 1 h at room temperature in a blocking solution containing 10% normal donkey serum and 0.3% triton X-100 in PBS and incubated overnight at 4°C with primary antibodies: goat anti-mCherry (1:1,000; SicGen), chicken anti-GFP (1:1000; Aves Labs #1020) and/or mouseIgG2A anti-ChR2 (1:200; ARP Inc). Sections were rinsed three times in PBS and incubated for 60–90 min at room temperature or overnight at 4°C with the Alexa Fluor 488- and 594 and 647-conjugated secondary antibodies (1:500; Thermo Fisher Science or Jackson ImmunoResearch).

### Rabies tracing

For tracing afferents from NDNF^+^ neurons in V1, stereotactic injections were performed between P30-P35 in the case of adult mice. AAV-helpers and N2C-RV-mCherry were diluted with PBS at a ratio of 1:1 and 50nl was injected using NanojectIII at 1nl/s in V1 (AP-3.5mm, ML-2.5mm, DV-0.20mm). Animals were perfused 14 days later. For developmental time points, stereotaxic injections were performed using a neonate adapter (Harvard apparatus). Mouse pups were anesthetized by hypothermia and stereotaxically injected with the viruses at P1 (From Lambda: AP+0.2mm, ML-1.60mm, DV-0.1mm). Animals were perfused 14 days later at P15. All coordinates were determined to target mainly L1 of the cortex. In the case of RV tracing from L4 neurons, AAV helpers and RV were injected at a depth of DV-0.5mm.

### Viruses

*For rabies tracing:* AAV2/1-DIO-helper virus encoding N2c-G-P2A-TVA-P2A-eGFP was expressed in a single AAV vector as described in (Pouchelon 2020). EnvA-pseudotyped CVS-N2C(ΔG)-FlpO-mCherry was used. The RabV CVS-N2c(DG)-mCherry-P2A-FlpO was a gift from Thomas Jessell (Addgene plasmid # 73471) and the N2C-RV were either produced, amplified and EnvA-pseudotyped in the lab, or generously shared by K. Ritola at Janelia Farms Research Center.

*Other viruses used in the paper:* AAV2/1-hSyn-hChR2(H134R)-EYFP was a gift from Karl Deisseroth (Addgene #26973); AAV2/1-hSyn-hM3D(Gq)-mCherry was a gift from Bryan Roth (Addgene #50474); AAV(PHP-eb)-hSyn-DIO-hM3D(Gq)-mCherry was a gift from Bryan Roth (Addgene #44361); AAV2/1-Ef1a-fDIO-hChr2-eYFP was a gift from Karl Deisseroth (Addgene 55639); pAAV2/1-EF1a-DIO-hChR2-eYFP was a gift from Karl Deisseroth (Addgene #20298); AAV2/1-hSyn-fDIO-DTA (NYUAD); AAV2/1-hSyn-hM4D(Gi)-tagBFP (NYUAD).

### Imaging analysis

Each brain section containing labelled cells was acquired as a tiled image on a motorized tiling scope Zeiss Axio Imager A1. Starter cells (colocalization of GFP^+^ AAV-helpers and mCherry^+^ RV) were manually quantified on ImageJ software. Brains with more than 5 non-L1 starter cells were discarded. mCherry^+^ retrogradely labeled cells were registered for each region of the Allen Reference Brain atlas for adult brain and of the “Atlas of Developing Mouse Brain at P6” from George Paxinos 2006. The total number of retrogradely labeled cells were normalized to the total number of cells labeled in the entire brain for the analysis in Fig. 1E; to the total number of retrogradely labeled cells within V1 (Fig. S1A-B) or to the total number of retrogradely labeled cells in the visual thalamus (Fig. S1C-D).

### Stereotaxic Injections for optogenetics and slice recording

Optogenetic stimulation of labeled axons was performed after expression of ChR2 (AAV1-hSyn-hChR2(H134R)-eYFP in the case of ACC injections [AP 0.5mm, ML 0.35mm, DV 0.5mm], or AAV1-EF1a-DIO-hChR2(H134R)-eYFP in the dLGN of Vipr2-IRES-Cre-D mice using stereotaxic injection at P1 for developmental time point [From Lambda: AP −0.8mm, ML 1.3mM, DV 1.4mm]. Either WT mice injected with AAV1-hSyn-hChR2(H134R)-eYFP or Vipr2-IRES-Cre-D mice were injected with AAV1-EF1a-DIO-hChR2(H134R)-eYFP at ∼P30-35 for the adult time point [AP −2.5mm, ML 2.5mm, DV 3mm]. No differences in the strength were found between the two cases.

### In vitro electrophysiology

P13-P15 were decapitated and the brain was quickly removed and immersed in ice-cold oxygenated (95% O2 / 5% CO2) sucrose cutting solution containing 87 mM NaCl, 2.5 mM KCl, 2 mM MgCl_2_, 1 mM CaCl_2_, 1.25 mM NaH_2_PO_4_, 26 mM NaHCO_3_, 10 mM glucose and 75 mM sucrose (pH 7.4). 300 μm thick coronal slices were cut using a Leica VT 1200S vibratome through V1. Slices were recovered in a holding chamber with artificial cerebrospinal fluid (aCSF) containing 124 mM NaCl, 20 mM Glucose, 3 mM KCl, 1.2 mM NaH_2_PO_4_, 26 mM NaHCO_3_, 2 mM CaCl_2_, 1 mM MgCl_2_ (pH 7.4) at 34 °C for 30 minutes and at room temperate for at least 45 minutes prior to recordings. For adult recordings, mice were perfused with NMDG cutting solution containing 92mM NMDG, 2.5mM KCl, 1.2mM NaH_2_PO_4_, 30mM NaHCO_3_, 20mM HEPES, 25mM glucose, 5mM sodium ascorbate, 3mM sodium pyruvate, 0.5mM CaCl_2_, 10mM MgSO_4_. During recovery, the NaCl was gradually added as described in (Ting, Lee et al. 2018). For recordings, slices were transferred to an upright microscope (Zeiss) with IR-DIC optics. Cells were visualized using a 40x water immersion objective. Slices were perfused with aCSF in a recording chamber at 2 ml/min at room temperature. All slice preparation and recording solutions were oxygenated with carbogen gas (95% O_2_, 5% CO_2_, pH 7.4). Patch electrodes (3–6 MΩ) were pulled from borosilicate glass (1.5 mm OD, Harvard Apparatus). For all recordings patch pipettes were filled with an internal solution containing: 125 mM Cs-gluconate, 2 mM CsCl, 10 mM HEPES, 1 mM EGTA, 4 mM MgATP, 0.3 mM NaGTP, 8 mM Phosphocreatine-Tris, 1 mM QX-314-Cl, equilibrated with CsOH at pH 7.3 or 130 K-Gluconate, 10 KCl, 10 HEPES, 0.2 EGTA, 4 MgATP, 0.3 NaGTP, 5 Phosphocreatine and 0.4% biocytin, equilibrated with KOH CO2 to a pH=7.3.

Recordings were performed using a Multiclamp 700B amplifier (Molecular Devices) and digitized using a Digidata 1440A and the Clampex 10 program suite (Molecular Devices). Voltage-clamp signals were filtered at 3 kHz and recorded with a sampling rate of 10 kHz. Recordings were performed at a holding potential of −70 mV. Cells were only accepted for analysis if the initial series resistance was less than 40 MΩ and did not change by more than 20% during the recording period. The series resistance was compensated at least ∼50% in voltage-clamp mode and no correction were made for the liquid junction potential. Whole-cell patch-clamp recordings were obtained from L1cINs and pyramidal-shaped neurons in L2/3 or L4 located in the same column. To activate afferents expressing hChR2, blue light was transmitted from a collimated LED (Mightex) attached to the epifluorescence port of the upright microscope. 5 ms pulses of a fixed light intensity were directed to the slice in the recording chamber via a mirror coupled to the 40x objective. Flashes were delivered every 15 s for a total of 10 trials. The LED output was driven by a transistor-transistor logic output from the Clampex software. In some cases, recordings were performed in the presence of 1 µm TTX and 1 mM 4-AP (Tocris) to reveal pure monosynaptic inputs. In the cases where cell identity needed to be determined, TTX and 4-AP were not used.

Passive and active membrane properties were recorded in current clamp mode by applying a series of hyperpolarizing and depolarizing current steps and the analysis was done in Clampfit (Molecular Devices). The cell input resistance was calculated from the peak of the voltage response to a 100 pA hyperpolarizing 1 second long current step according to Ohm’s law. Analysis of the action potential properties was done on the first spike observed during a series of depolarizing steps. Threshold was defined as the voltage at the point when the slope first exceeds a value of 20 Vs^-1^. Rheobase was defined as the amplitude of the first depolarizing current step at which firing was observed. Analysis of spontaneous inhibitory events was done using Clampfit’s threshold search.

Data analysis was performed off-line using the Clampfit module of pClamp (Molecular Devices) and Prism 8 (GraphPad). The amplitude of evoked synaptic currents was obtained by averaging the peak amplitude of individual waveforms over 10 trials per cell. EPSC amplitudes recorded from L1 cINs were then normalized for injection size by dividing the average EPSC by the evoked current amplitude from a putative L2/3 or L4 pyramidal neuron in the same column of each slice.

### CNO injections

CNO (diluted in DMSO and saline) was injected at a concentration of 0.01mg/kg of body weight and was injected daily starting at P5. Injections were made in the milk sac of the pups, and later injections were made intraperitoneally.

### Neurolucida tracing

Sections containing the axons of interest were imaged on a Zeiss LSM 800. Z-stacks of images were then loaded into Neurolucida 360 (MBF Biosciences) and trees were reconstructed using the ‘user guided’ option with Directional Kernels.

### Statistical Analysis

No prior test for determining sample size was conducted. All statistical analyses were performed using Prism (GraphPad). Statistical significance was tested with non-paired, two-sided t-test, with a 95% confidence interval or One-Way ANOVA with Bonferroni correction for multiple comparisons.

## ACKNOWLEDGEMENTS

We thank Dr. Anne Takesian and Dr. Chinfei Chen for their valuable comments on the manuscript. We also thank Dr. Tim Burbridge for demonstrating retinal injections. This work was supported by a Goldenson Foundation and a Hearst foundation grant (to LAI) and by grants from the National Institutes of Health (NIH): MH071679, NS08297, NS074972, MH095147 to GF and grants P01NS074972, R01NS107257, and R01NS110079 to BR, as well as support from the Simons Foundation (SFARI) (to G.F).

## AUTHOR CONTRIBUTIONS

LAI and GF conceived the project and wrote the manuscript. LAI carried out injections, histology and recording experiments. SH carried out dLGN-V1 recording experiments and assisted with dLGN virus injections. MFO and SV assisted with histology experiments. MS assisted with rabies quantification and neurolucida reconstructions. QX produced some of the viruses used in the study. RM and BR contributed to the design of the experiments and discussion.

## COMPETING INTERESTS

The authors declare no competing interests.

## Notes

### Competing Interest Statement

The authors have declared no competing interest.

### Summary of Updates

Figures with a margin on top so that the stamper doesnt overlap on the figure itself.

