## Supplementary Figures for "Bottom-up inputs are required for the establishment of top-down connectivity onto cortical layer 1 neurogliaform cells"

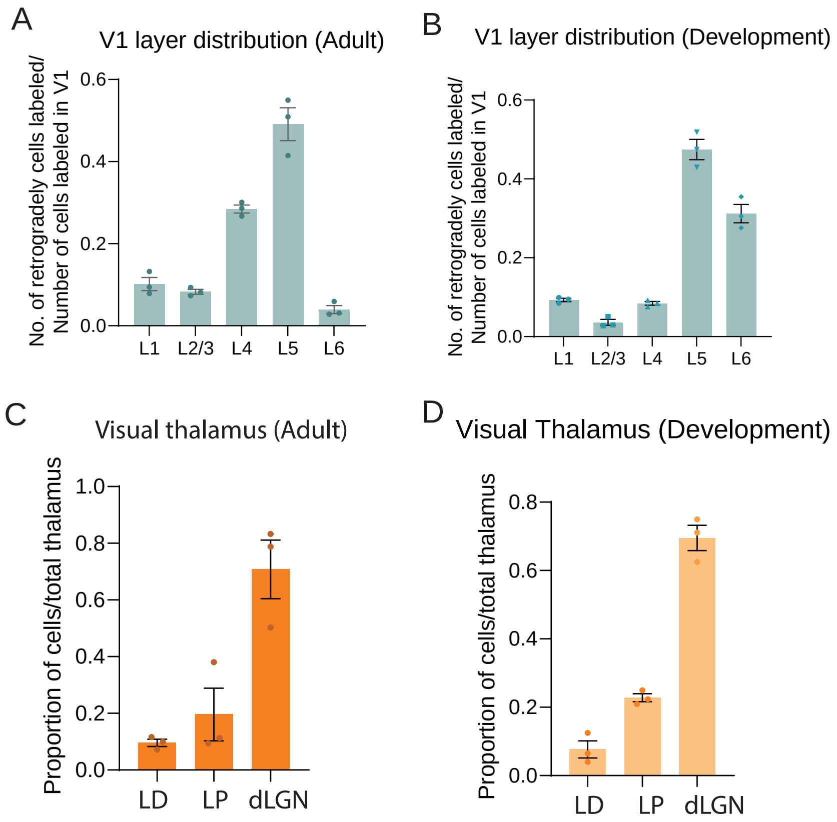


**Figure S1. Distribution of RV-retrogradely labeled cells from NDNF+ve L1 cINs. Related to Figure 1**

(A) Local input V1 layer distribution in adults

(B) Local input V1 layer distribution in development

(C) Visual thalamic nuclei inputs in adult

(D) Visual thalamic nuclei inputs in development

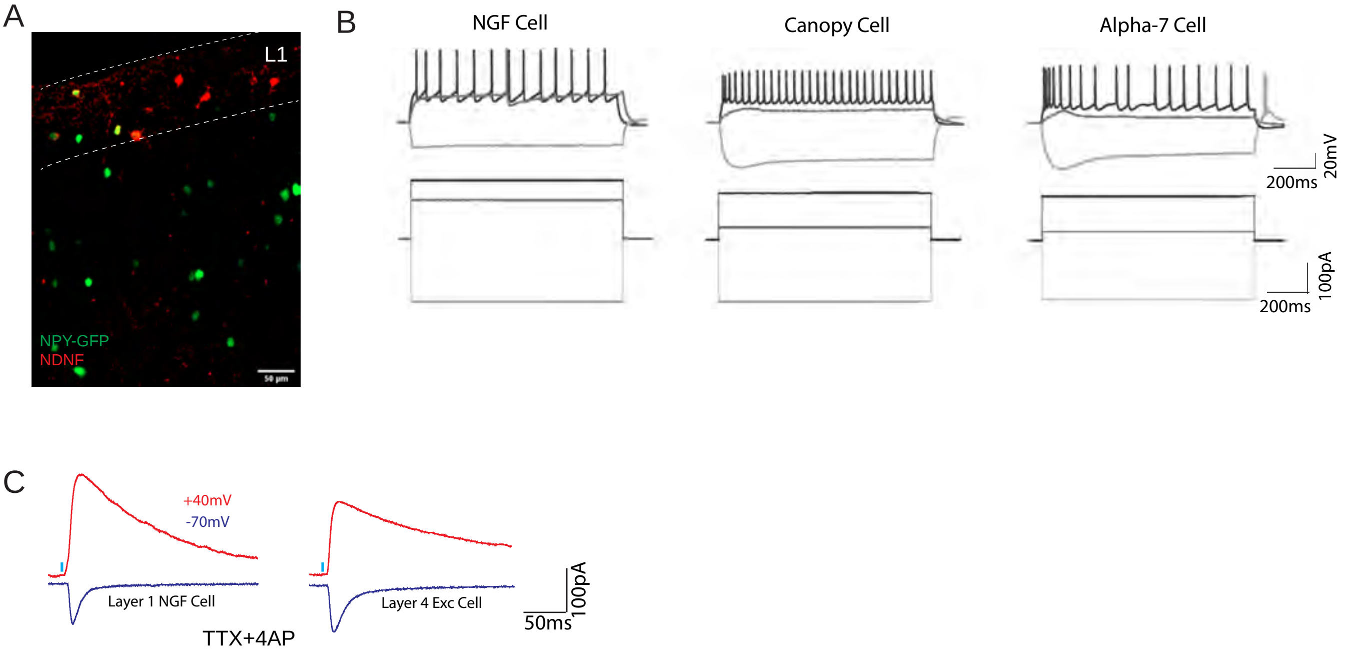


**Figure S2. Identification and intrinsic responses of the different L1 cIN subtypes during development . Related to Figure 2**

(A) Image showing distribution of NGF and canopy cells in L1 of V1. Scale Bar=50μm

(B) Example traces from the three major L1 cIN subtypes, NGF cell, Canopy cell and alpha-7 expressing cell.

(C) Responses of a L1 NGF cell and L4 excitatory neuron in the same column to optogenetic stimulation of dLGN fibers during development. We confirmed that these connections were monosynaptic and had an NMDA component using Cs+ internal solution together with TTX and 4AP. As previously described (De Marco et. al 2015), we found that the NMDA current in L1 cINs was larger than that seen in L4 excitatory cells

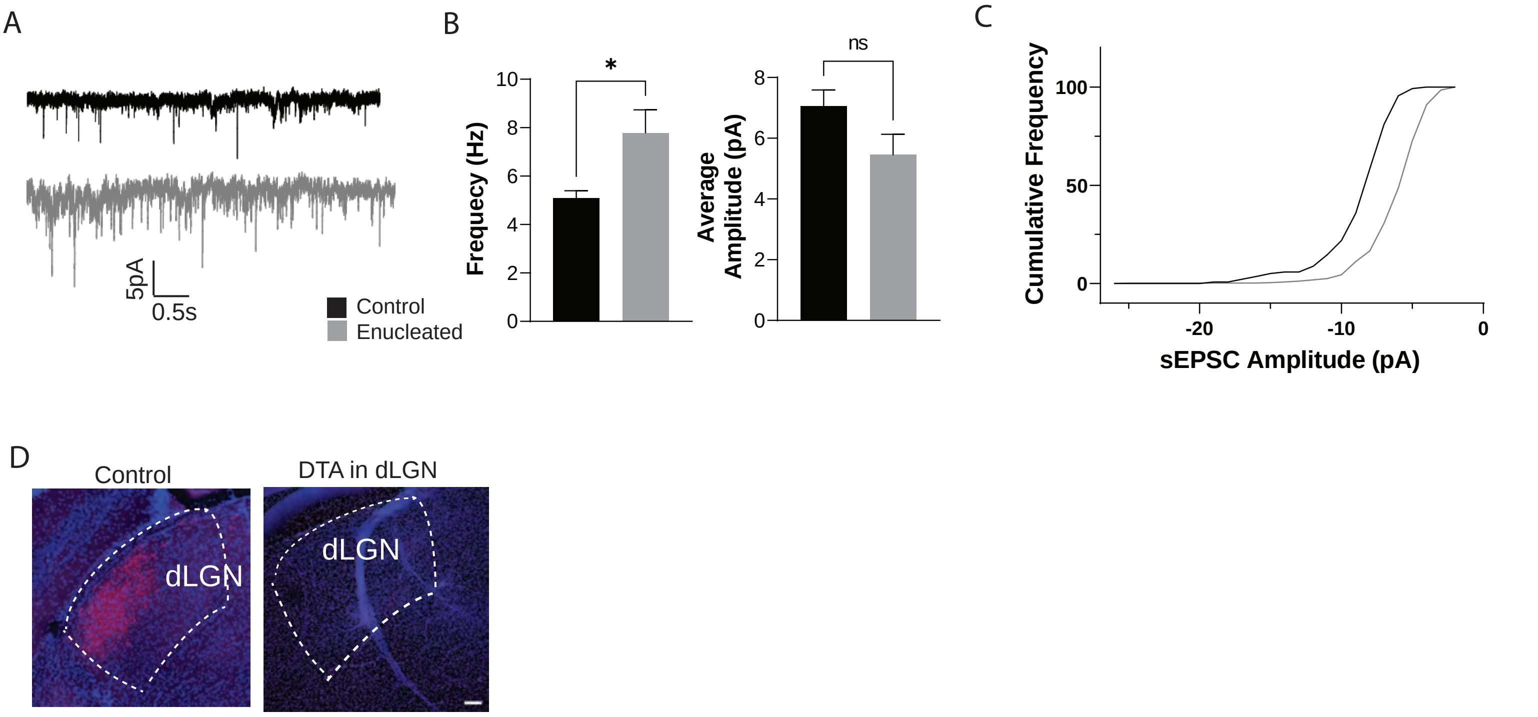


Figure S3. Related to Figure 4

(A) Example spontaneous currents from L1 cINs in either control (top) or enucleated animals (bottom).

(B) Frequency and ampliude of the spontaneous events recorded from all L1 cINs under these conditions. *=<0.05

(C) Cumulative frequency distribution of amplitudes

(D) Images demonstrating the DTA strategy is effective in ablating retrogradely labeled dLGN neurons. Scale Bar=100μm

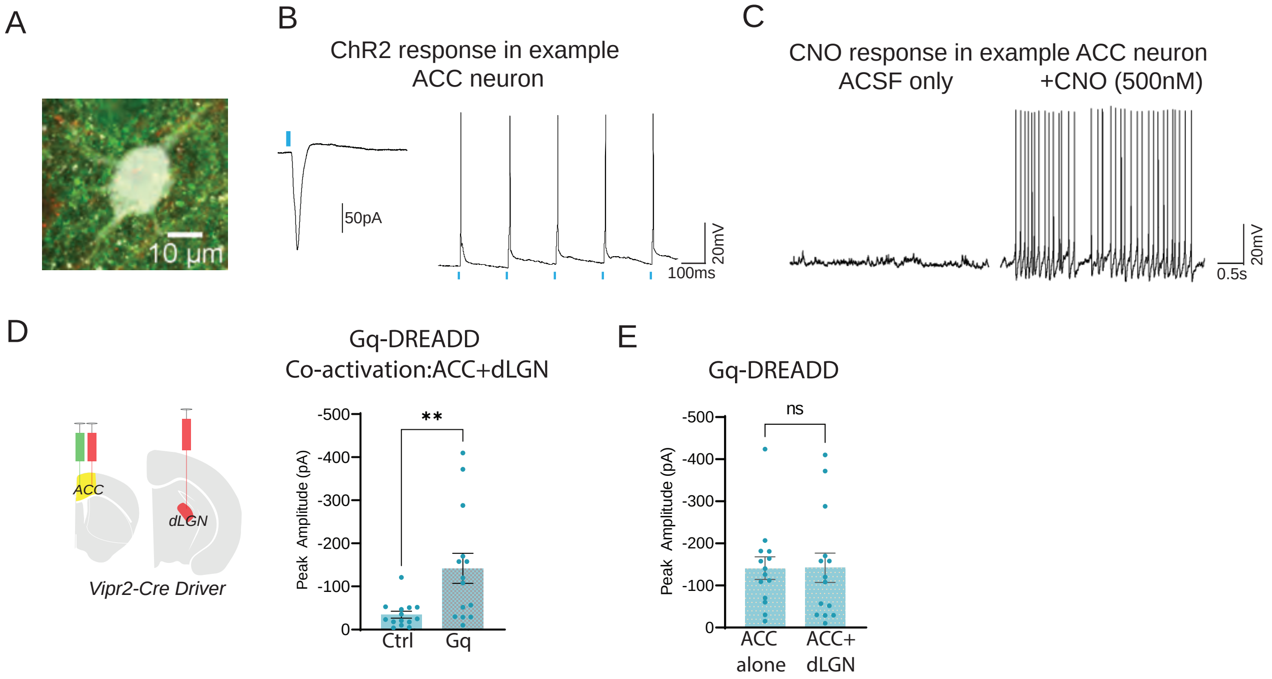


**Figure S4. Gq DREADD activation. Related to Figure 5**

(A) Biocytin filled neuron recorded from the ACC expressing both eYFP and mCherry

(B) Response of neuron in A to optogenetic stimulation; voltage clamp recording to a single 5ms pulse stimulation (left panel), current clamp recording to a 5Hz stimulation (right panel).

(C) Spontaneous response (under current clamp recording) of the same neuron to ACSF (left panel) or bath application of CNO (right panel)

(D) Schematic of the co-incident activation of bottom-up and top-down pathway (left panel). Peak amplitude responses of L1 cIN to optogenetic stimulation of ACC axons in V1 under control or CNO conditions

(E) Comparison of NGF cell responses when only ACC was over-activated compared to when both dLGN and ACC were simultaneously activated
